# Crystal structure of the TbBILBO1 N-terminal domain reveals a ubiquitin fold with a long rigid loop for the binding of its partner

**DOI:** 10.1101/738153

**Authors:** Keni Vidilaseris, Nicolas Landrein, Yulia Pivovarova, Johannes Lesigang, Niran Aeksiri, Derrick R. Robinson, Melanie Bonhivers, Gang Dong

## Abstract

BILBO1 was the first characterized component of the flagellar pocket collar (FPC) in trypanosomes. The N-terminal domain (NTD) of BILBO1 plays an essential role in *Trypanosoma brucei* FPC biogenesis and is thus vital for the parasite’s survival. Here we report a 1.6-Å resolution crystal structure of TbBILBO1-NTD, which revealed a conserved horseshoe-like hydrophobic pocket formed by an unusually long loop. Mutagenesis studies suggested that another FPC protein, FPC4, interacts with TbBILBO1 via mainly contacting the three conserved aromatic residues W71, Y87 and F89 at the center of this pocket. Overall, we have determined the binding site of TbFPC4 on TbBILBO1-NTD, which may provide a basis for rational drug design in the future.

## INTRODUCTION

*Trypanosoma brucei* is a protist parasite causing sleeping sickness and Nagana in Sub-Saharan Africa. At the base of its single flagellum is a bulb-like invagination of the plasma membrane called the flagellar pocket (FP), which is responsible for all endo-/exocytosis in the cell (1–3). Around the neck of the FP on its cytoplasmic face is an electron-dense structure called the flagellar pocket collar (FPC), which is essential for FP biogenesis and thus for the survival of the parasite (4).

TbBILBO1 was the first reported protein component of the FPC (5). It consists of four structural domains: a globular N-terminal domain; two central EF-hand motifs; a long coiled-coil domain; a C-terminal leucine zipper (6). Analysis using electron microscopy showed that TbBILBO1 forms long filaments *in vitro* (6,7). Based on its modular architecture and intrinsic property to form spiral-like structures when ectopically expressed *in vivo*, TbBILBO1 was proposed to act as the structural core to recruit other components during FPC assembly (8,9). Another confirmed FPC component besides BILBO1 is FPC4, which is a microtubule binding protein that interacts with microtubule through its N-terminal part and with TbBILBO1-NTD via its C-terminal region (8). However, how TbFPC4 binds to TbBILBO1 remains unclear.

We have previously reported an NMR structure of TbBILBO1-NTD (aa1-110) and identified a conserved surface patch essential for the function of TbBILBO1 in the parasite (10). However, due to the limitation of the method, that structure had a relatively low resolution, particular for the ill-defined surface patch because of the lack of assigned signals of the C-terminal loop. It was also unclear to us at that time what the binding partner of this conserved patch was.

Here we report a 1.6-Å resolution crystal structure of TbBILBO1-NTD, which, with a longer sequence in the final structure (aa1-115), revealed a unique horseshoe-like pocket harboring multiple conserved aromatic residues. This pocket is formed by an extended loop at the C-terminus of TbBILBO1-NTD, spanning residues 96-115. Structure-based mutagenesis studies suggested that TbFPC4 contacts primarily the three conserved aromatic residues in the center of the horseshoe-shaped pocket. The binding interface also extends to the peripheral area close by the gap of the horseshoe-like the pocket. Overall, our work unveils a unique structural feature of TbBILBO1-NTD, including a long rigid loop and a conserved horseshoe-like aromatic pocket, and sheds light on how TbFPC4 is docked at the pocket to achieve high binding affinity.

## EXPERIMENTAL PROCEDURES

### Cloning and site-directed mutagenesis

Cloning of full-length (aa1-587) and NTD-EFh (aa1-249) constructs of TbBILBO1 has been reported previously, where a stop codon was added at the 3’ end of the cloned sequence coding for the whole protein or residues 1-249 (10,11). For mutagenesis studies, a new TbBILBO1-NTD construct based on the crystal structure, which contains residues 1-120, was cloned into the expression vector pET15b (Novogen) between *Nde*I and *BamH*I sites. The resulting construct provides an N-terminal 6×His tag, which is cleavable by the thrombin protease.

For TbFPC4, we cloned the sequence encoding its C-terminal domain (CTD, aa357-404), which was reported to be sufficient for TbBILBO1 binding (8), from genomic DNA into a custom vector SUMO15b. The vector provides an N-terminal 6×His-SUMO tag that is cleavable by the Sentrin-specific protease 2 (SENP2) protease.

Two single-residue mutants of TbBILBO1-NTD, Y64A and W71A, were generated by site-directed mutagenesis using a QuikChange kit (Stratagene) according to the manufacturer’s instructions. Incorporation of mutations was confirmed by DNA sequencing.

### Protein expression and purification

Recombinant TbBILBO1-NTD and TbFPC4-CTD proteins were expressed in *Escherichia coli* (strain BL21-DE3). Briefly, the cloned constructs for protein expression were used to transform competent bacterial cells. The cells were grown in Luria-Bertani (LB) medium at 37°C to an OD_600_ of 0.6-0.8, and then subjected to cold shock on ice for 30 min. Protein expression was induced by addition of 0.5 mM isopropyl β-D-1-thiogalactopyranoside (IPTG), and cultures were further incubated at 16°C overnight (14-16 h). Cells were harvested by centrifugation in a Sorvall GS3 rotor (6,000×g, 12 min, 4°C) and then resuspended in 20 ml of lysis buffer (20 mM Tris-HCl pH 8.0, 100 mM NaCl, 20 mM imidazole, 5% (v/v) glycerol) per L of cell culture.

Bacteria were lysed in an EmulsiFlex-C3 homogenizer (Avestin) and cell debris was pelleted by centrifugation (40,000×g, 40 min, 4°C). The supernatant was filtered (0.45-μm pore size, Amicon) and loaded onto a Ni-HiTrap column (GE Healthcare) pre-equilibrated with the same lysis buffer. The column with bound proteins was washed with 5 × column volume (cv) of lysis buffer, and bound protein was eluted using a linear gradient concentration of imidazole (20 -500 mM, 20×cv) in the same lysis buffer.

The N-terminal 6×His tag of TbBILBO1-NTD and the 6×His-SUMO tag of TbFPC4-CTD were cleaved off by incubating the pooled fractions of interest with ~2% (w/w) of thrombin and ~1% (w/w) of SENP2, respectively (4°C, overnight). Target proteins were further purified on a Superdex-200 16/60 column (GE Healthcare) pre-equilibrated with a running buffer containing 20 mM Tris-HCl (pH 8.0) and 100 mM NaCl. The eluted proteins were pooled and used for subsequent binding tests or native gel electrophoresis. Protein concentration was determined using Ultraviolet (UV) absorbance at 280 nm. The molar concentration of both proteins was calculated according to the UV-based concentration measurement (mg/ml) and the extinction coefficients (16960 M^−1^·cm^−1^ and 8480 M^−1^·cm^−1^ for TbBILBO1-NTD and TbFPC4-CTD, respectively) obtained using the ExPASy Server (12).

### Crystallization and structure determination

Crystallization of TbBILBO1-NTD and diffraction data collection have been reported previously (11). To briefly sum it up, following purification on Ni-HiTrap column selenium-methionine (SeMet) substituted TbBILBO1-NTD-EFh was incubated with thrombin to release the NTD (there is a thrombin cleavage site after residue R175). After further purification by ion exchange and size exclusion chromatography, protein sample corresponding to residues 1-175 was used to set up crystallization trials. The resulting plate-like crystals were used to collect a highly redundant single anomalous dispersion (SAD) dataset at the absorption edge of selenium (Se, λ = 0.9792 Å) on the ID14-4 beamline (ESRF). Structure determination was carried out using the SAD method exploiting the three selenium (Se) sites in the SeMet-substituted methionine residues (M1, M63, and M107). All Se sites were ordered and could be readily located, and experimental maps were calculated using AutoSol in the software suite Phenix (13), which built the majority of the structural model automatically. The model was further manually checked and rebuilt (for loops and terminal residues) using the program COOT (14). Structure refinement was carried out using phenix.refine (13).

### Isothermal titration calorimetry (ITC)

ITC measurements were performed on a MicroCal™ iTC200 microcalorimeter (GE Healthcare). Protein samples of TbBILBO1-NTD and TbFPC4-CTD were dialyzed overnight against 100 × volume of the binding buffer containing 20 mM Tris-HCl (pH 8.0) and 100 mM NaCl. A typical ITC titration experiment consisted of 20 injections of TbFPC4 (300 μM), first with 1 × 0.2 μl and then with 19 × 2 μl, into a reaction cell filled with 200 μl of TbBILBO1-NTD (30 μM).

All ITC measurements were carried out under constant stirring at 350 rpm, and each injection lasted for 4 s with a 180-s interval between two injections. Titration peaks were analyzed using the Origin 7.0 Microcal software and corrected for FPC4-CTD dilution heat measured by injecting the protein into only the buffer using the same protocol described above. Non-linear least squares fitting using one binding site model was used to calculate the association constant (Ka) and stoichiometry values. Dissociation constants (Kd) were calculated according to the formula Kd = 1/Ka. Each of the reported ITC results is the representative one of at least three independent measurements.

### Static light scattering (SLS)

SLS measurements were carried out by coupling size-exclusion chromatography (SEC) with mass determination. In each measurement, a 50-μl protein sample (1-2 mg/ml) was analyzed on a Superdex S-200 10/300 GL column (GE Healthcare) pre-equilibrated with a buffer containing 20 mM Tris-HCl (pH 8.0), 100 mM NaCl, 1 mM dithiothreitol, and 1% (v/v) glycerol. Data analyses were carried out using the ASTRA software provided by the manufacturer.

### Electrophoretic mobility shift assay (EMSA)

All EMSA experiments were carried out on 5% (w/v) native polyacrylamide gels using Tris-acetate-EDTA (TAE) buffer containing 40 mM Tris, 20 mM acetic acid, and 1 mM EDTA. The gels were run at 150 V for 2 h at 4°C. The TbBILBO1-NTD/TbFPC4-CTD complex purified by SEC was loaded directly on the native gel in the presence of 10% (v/v) glycerol to allow the samples to sediment within the wells. In all other binding tests, the separately purified protein samples (~1 mg/ml) were mixed in a 1:1 ratio and incubated overnight at 4°C before loading onto the native gel. Single protein samples with the same concentration as the complexes were loaded on the same gel as controls.

### T. brucei cell culture and mutant protein production

The *in vivo* work in the parasite described in this study used the procyclic (PCF) *T. brucei* 427 29.13 co-expressing the T7 RNA polymerase and the tetracycline repressor (15,16). Cells were cultured at 27°C in SDM79 medium (Sigma-Aldrich) containing 10% (v/v) heat-inactivated fetal calf serum, hemin 20 μg/ml, hygromycin 25 μg/ml, neomycin 10 μg/ml. Phleomycin (5 μg/ml) was also added for the cell lines expressing the Ty1-tagged TbBILBO1 constructs (10). 3 × 10^7^ cells were transfected with 10 μg of *Not*I linearized plasmids using the AMAXA electroporator (Lonza, program X-001) as described (15), with the transfection buffer (17). Ty1-tagged TbBILBO1 mutant cell lines – two single mutants Y64A and W71A, and a triple mutant D65K/E66K/E67K (DEE/KKK) - were generated by site-directed mutagenesis (Quick Change Lightning Site directed Mutagenesis Agilent Kit 210518) using the pLew100-TbBILBO1 plasmid and transfection of the Tb427 29.13 cell line (10). Incorporation of mutations was confirmed by sequencing of the plasmids and of genomic DNA after cell line selection. Expression of Ty1-tagged WT-TbBILBO1 and mutant forms was induced by adding 20 ng/ml of tetracycline in the culture medium.

### Mammalian cell culture

U-2 OS cells (human bone osteosarcoma epithelial cells, ATCC Number: HTB-96 (18)) were grown in D-MEM Glutamax (Gibco) supplemented with final concentrations of 10% (w/v) fetal calf serum (Invitrogen), 100 units/ml of Penicillin (Invitrogen), and 100 μg/ml of Streptomycin (Invitrogen) at 37°C plus 5% (w/v) CO_2_. Exponentially growing U-2 OS cells in 24-well plates with glass coverslips were lipotransfected as described in (19) with 0.5μg of DNA using Lipofectamine 2000 in OPTIMEM (Invitrogen) according to the manufacturer’s instructions, and processed for immunofluorescence imaging 24 hours post-transfection. To transiently express TbBILBO1 and its mutants in the heterologous mammalian system together with TbFPC4-GFP, we generated the pcDNA3-TbBILBO1-Y64A and pcDNA3-TbBILBO1-W71A mutant plasmids using the pcDNA3-TbBILBO1 plasmid (9) and site-directed mutagenesis as described above.

### Immunofluorescence

Immunofluorescence on detergent-extracted U-2 OS cells was done as described in (8).

### Western blotting

Trypanosome whole cell protein lysates (2 × 10^6^ cells per well) were separated using SDS-PAGE (10% gels) and transferred by semi-dry blotting (BioRad) for 45 min at 25 V to PVDF membranes. After a 1-h blocking step in 5% skimmed milk in TBS, 0.2% Tween-20 (blocking solution BS), the membranes were incubated with the primary antibodies diluted in BS – anti-Ty1 (BB2 mouse monoclonal, 1:10,000 (20)) and anti-Enolase (loading control, rabbit polyclonal, 1:10,000 (21)). After one wash in BS, one wash in 1M NaCl and one wash in BS, the membranes were incubated with HRP-coupled secondary antibodies diluted 1:10,000 in BS (anti-mouse IgG Jackson 515-035-062, anti-rabbit IgG Sigma A-9169), washed twice in BS and twice in TBS, and visualized using the Clarity Western ECL Substrate kit (Bio-Rad) with an ImageQuant LAS4000. After stripping (two washes of 5 min in 100 mM glycine-HCl pH 2.3, 1% (w/v) SDS, 0.1% (w/v) NP40), washing in TBS then blocking in BS, the membranes were incubated with a rabbit polyclonal anti-TbBILBO1 (rabbit polyclonal (8)) and imaged as described above.

### Accession code

Coordinates and structure factors of the TbBILBO1-NTD crystal structure have been deposited in the Protein Data Bank (PDB) under accession code 6SJQ.

## RESULTS

### 1.6-Å resolution crystal structure of TbBILBO1-NTD

As reported previously, TbBILBO1-NTD-EFh was expressed in bacteria and purified on three consecutive chromatographic columns to obtain highly homogeneous sample (11). The thrombin-cleaved product of SeMet-substituted TbBILBO1-NTD-EFh, corresponding to residues 1-175, was crystallized after initial screening and subsequent optimization. The resulting crystals belonged to space group P2_1_ (*a* = 29.69 Å, *b* = 50.80 Å, *c* = 37.22 Å; β = 94.61°). The structure was solved to 1.6-Å resolution employing the SAD method using diffraction data collected at the absorption edge of selenium (Table 1). The final structure contained residues 2-115 of TbBILBO1-NTD, together with 135 bound water molecules that were confidently modeled on the basis of significant electron densities and satisfied hydrogen bonding interactions. The refined model has *R*_work_ and *R*_free_ of 16.9% and 19.8%, respectively.

The crystal structure revealed that TbBILBO1-NTD consists of five β strands (β1 - β5) and a single long α helix (Fig. 1A). The 2Fo-Fc electron density map is excellent for all residues in the structure as well as the bound water molecules (Fig. 1B). Notably, at this resolution one can easily spot the holes in the electron density of all aromatic rings as well as the proline residues. The first two β strands, β1 & β2, are much longer than β3-β5 and provide the majority of the contacting interface with the helix. These five β strands together form a slightly twisted β sheet, with the α-helix packed diagonally along the back side of the sheet (Fig. 1C). Overall the structure is topologically the same as the canonical ubiquitin fold, similarly to the previously reported NMR structure (10).

**FIGURE 1.**
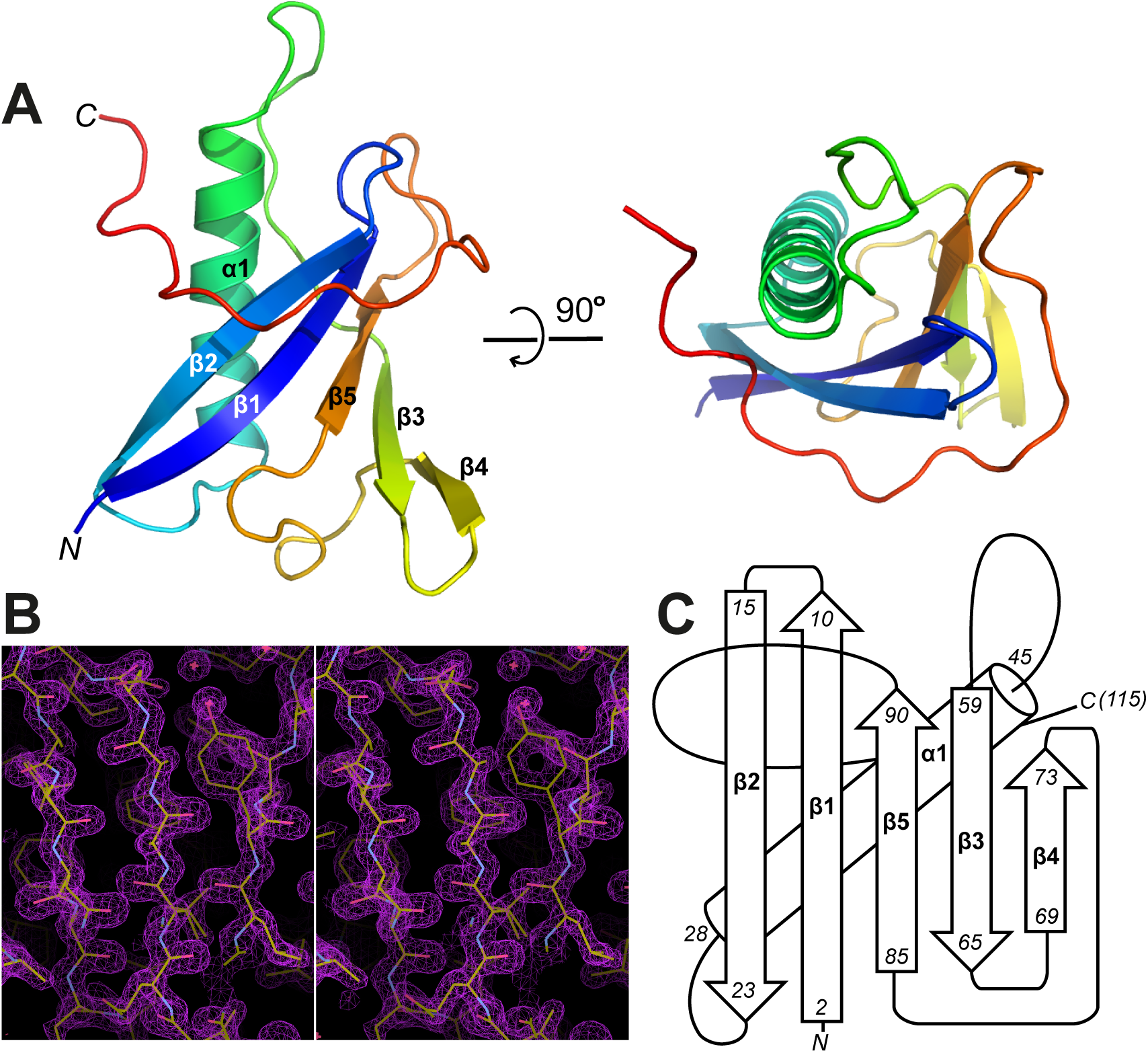
Crystal structure of TbBILBO1-NTD. *A*, Ribbon diagram of the TbBILBO1-NTD structure with two orthogonal views. The structure is color-ramped from blue at the N-terminus to red at the C-terminus. The five β strands (β1 - β5) and the single α helix (α1) are labeled. *B*, Stereo view of a part of the 2Fo-Fc map contoured at 2 σ level. *C*, Secondary structure diagram of TbBILBO1-NTD, with the residue ranges for each structural elements indicated.

### The C-terminal tail of TbBILBO1-NTD forms an extended loop

The crystal structure of TbBILBO1-NTD is overall similar to the previously reported NMR structure. The average root-mean-square deviation (r.m.s.d.) between the two structures for all atoms within residues 1-95 is ~2.7 Å, and for only the ordered structures (i.e. α1 and β1-β5) is ~1.9 Å. However, the C-terminal extension (aa101-110) of TbBILBO1-NTD was poorly defined in the NMR structure due to the missing long-range nuclear Overhauser effect (NOE) signals for the residues in that region, and was previously proposed to be structurally dynamic (Fig. 2A). Interestingly, the crystal structure showed a rigid loop that is unusually long, spanning residues 96-115, and wrapping around β1, β2 and the C-terminal part of the α helix (Fig. 2B). This loop is structurally rigid as demonstrated both by the clear electron densities of all residues in the loop and by their comparable temperature factors to those of the residues in the connecting loops in the core structure (Fig. 2B and C). Interaction of the residues in the loop with the core structure is stabilized by more than 20 hydrogen bonds, either directly or mediated by water molecules, and many hydrophobic interactions (Fig. 2D).

**FIGURE 2.**
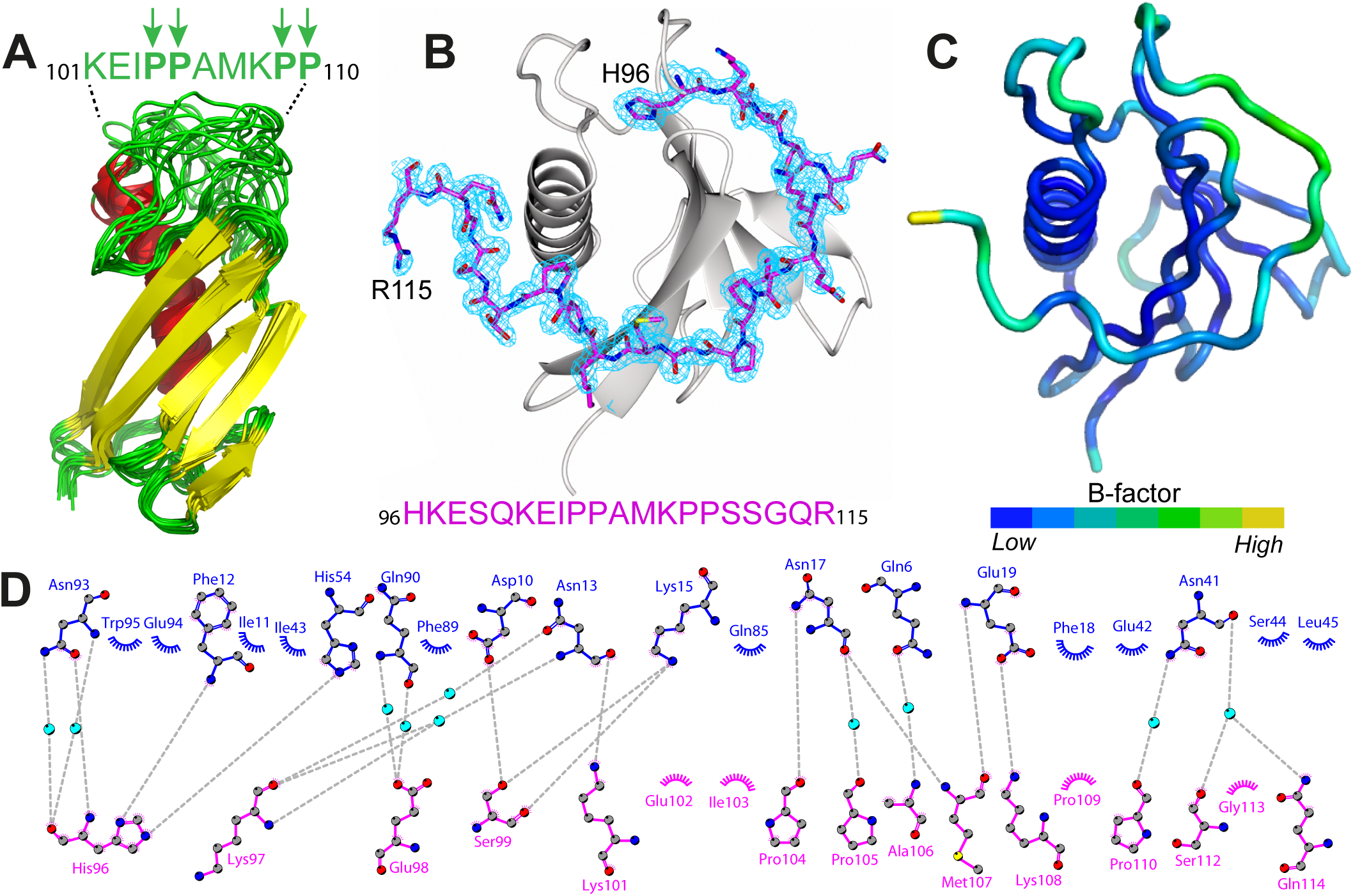
A long rigid loop is present at the C-terminus of TbBILBO1-NTD. *A*, Superimposed 10 energy-minimized NMR conformers of TbBILBO1-NTD. Residues in the dynamic C-terminal region (residues 101-110) are listed, with the four proline residues marked by arrows. *B*, Ribbon diagram of the crystal structure of TbBILBO1-NTD. The last twenty residues (aa96-115) comprising the loop are shown as sticks together with 2*F*_*o*_ - *F*_*c*_ electron density map around the residues contoured at 1.5 σ level. The plot was generated using CCP4mg (24). *C*, Ribbon diagram in the same orientation as in *B* and colored by temperature factors (B-factors), with blue being the most rigid and yellow the most dynamic. The average B-factor for the structure is 23 Å^2^. *D*, Details of interactions between the C-terminal tail (magenta) and the TbBILBO1-NTD core (blue). The plot was generated using DIMPLOT in the LigPlot plus suite (25). Residues involved in hydrogen bond formation are shown as ball-and-stick, with oxygen, nitrogen and carbon atoms colored in red, blue, and gray, respectively. Water molecules mediating inter-molecular hydrogen bond formation are shown as cyan-colored spheres. Grey dotted lines indicate hydrogen bonds. Non-bonded residues involved in hydrophobic interactions are shown as spoked arcs.

### TbBILBO1-NTD adopts a ubiquitin-like fold

To identify similar structures, we carried out heuristic PDB search on the Dali server (22) using the TbBILBO1-NTD structure as a query to search against all depositions in the Protein Data Bank (23). The reported match correlation matrix shows that the first 91 residues in the TbBILBO1-NTD structure forms a canonical ubiquitin-like fold, whereas the C-terminal extension from residue P92 to R115 forms a unique curved loop that is absent in all other known ubiquitin-like structures (Supplementary Fig. 1A). We chose the 11 top ranking structures, which have a Z-score equal or larger than 7.0 and r.m.s.d. ≤ 2.3 Å, to directly compare their structural similarity and difference with TbBILBO1-NTD. Superposition of these structures with the TbBILBO1-NTD structure shows that, while TbBILBO1-NTD shares a nearly identical core with other ubiquitin-like structures, its C-terminal loop is unprecedented and distinct (Supplementary Fig. 1B).

### The rigid conformation of the loop is not a crystallization artefact

Given the distinct conformation of the loop in the crystal structure, we check whether it is stabilized by extensive crystal contacts. We found that the loop is located in a cavity in the crystal, with the side-chains of most of the loop residues making no contact to neighboring molecules (Supplementary Fig. 2A). The 2Fo-Fc map also shows that, except for the C-terminal part, the rest of the loop interacts only with the core structure of the same molecule, but not with any other molecules in the neighborhood (Supplementary Fig. 2B). Overall, we can conclude that the conformation of the loop is in its native state rather than an artefact caused by crystal packing.

### The C-terminal loop helps to generate a hydrophobic horseshoe-shaped pocket

In the crystal structure, there is a sharp kink at the beginning of the long loop to bend the loop from the longitudinally orientated C-terminus of β5 (relative to the long axis of the β sheet) to a horizontally arranged conformation. At the kink point is the highly conserved residue H96, whose side chain is completely buried in a deep pocket (Fig. 3A). This rigid, curved loop generates a horseshoe-like pocket, with one side of the pocket being shaped completely by the loop (Fig. 3B). This pocket was not well defined in the previously reported NMR structure, mostly due to the poorly built loop that was detached from the rest of the structure (Fig. 3C). The central part of the pocket contains three aromatic residues, W71, Y87 and F89, with another aromatic residue Y64 in the proximity of the pocket (Fig. 3D). There are also two positive charged residues, K60 and K62, at one side of the pocket, and residue Q100 from the loop on the opposite side of the pocket (Fig. 3E).

**FIGURE 3.**
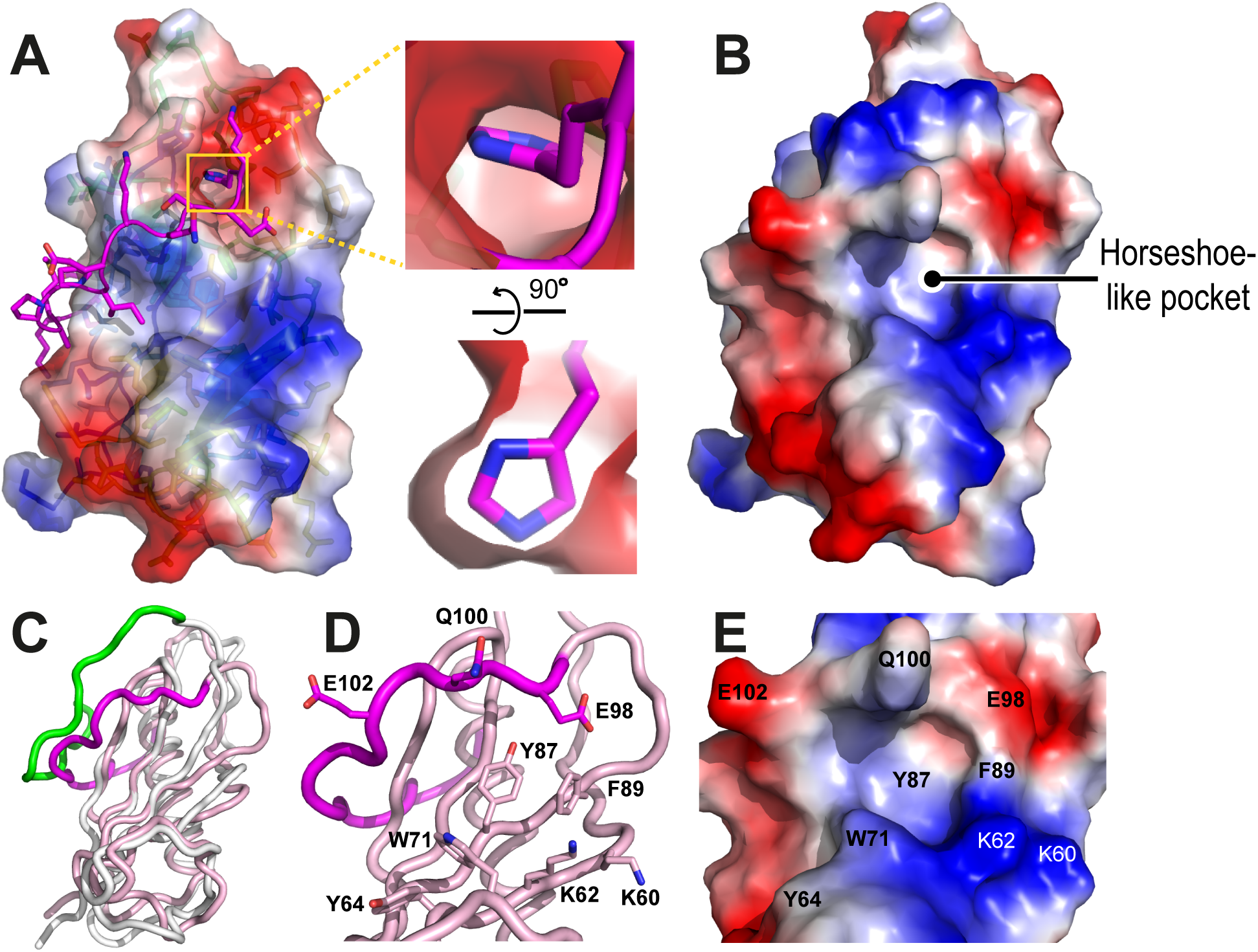
TbBILBO1-NTD has a conserved horseshoe-like surface pocket. *A*, Crystal structure of TbBILBO1-NTD with the C-terminal loop depicted as sticks and the rest of the structure as an electrostatic surface plot. Shown on the right are two orthogonal zoom-in views of the deeply buried side chain of residue H96 located at the beginning of the C-terminal loop. *B*, Full electrostatic plot with the same orientation as in *A* to show the horseshoe-like hydrophobic pocket. *C*, Superposition of the crystal structure (pink) onto the previously reported NMR structure (gray, PDB code: 2MEK). The corresponding C-terminal loops in the two structures are colored in magenta and green, respectively. *D*, Cartoon view of the crystal structure around the hydrophobic pocket of TbBILBO1-NTD, with residues in and around the pocket shown in sticks. *E*, Electrostatic plot of the pocket with the same orientation as in *D*.

### TbFPC4 binds to TbBILBO1-NTD mainly by contacting three central aromatic residues in the pocket

Previous studies have shown that several residues near the hydrophobic pocket are essential for the function of TbBILBO1 (10). Later studies demonstrated that TbFPC4 binds to a similar region of TbBILBO1-NTD (8). Based on these studies and the newly determined crystal structure of TbBILBO1-NTD, we used structure-based mutagenesis to define how TbFPC4 is orientated on TbBILBO1 in the formed complex.

We chose four sets of residues to test based on their high conservation among kinetoplastids and location within or close to the pocket. The two double mutants within the pocket, Y87A/F89A and K60A/K62A, have been previously tested for their influence on TbFPC4 binding (8). Based on the unique horseshoe-shaped conformation of the pocket, we further separately mutated two aromatic residues, including the bulky residue (W71) gating the gap of the horseshoe, and Y64, the aromatic residue just outside of the pocket (Fig. 4A). Each of the two residues were mutated to alanine to remove the bulky side-chains. We then checked how these mutants affect the interaction of TbBILBO1-NTD with TbFPC4.

**FIGURE 4.**
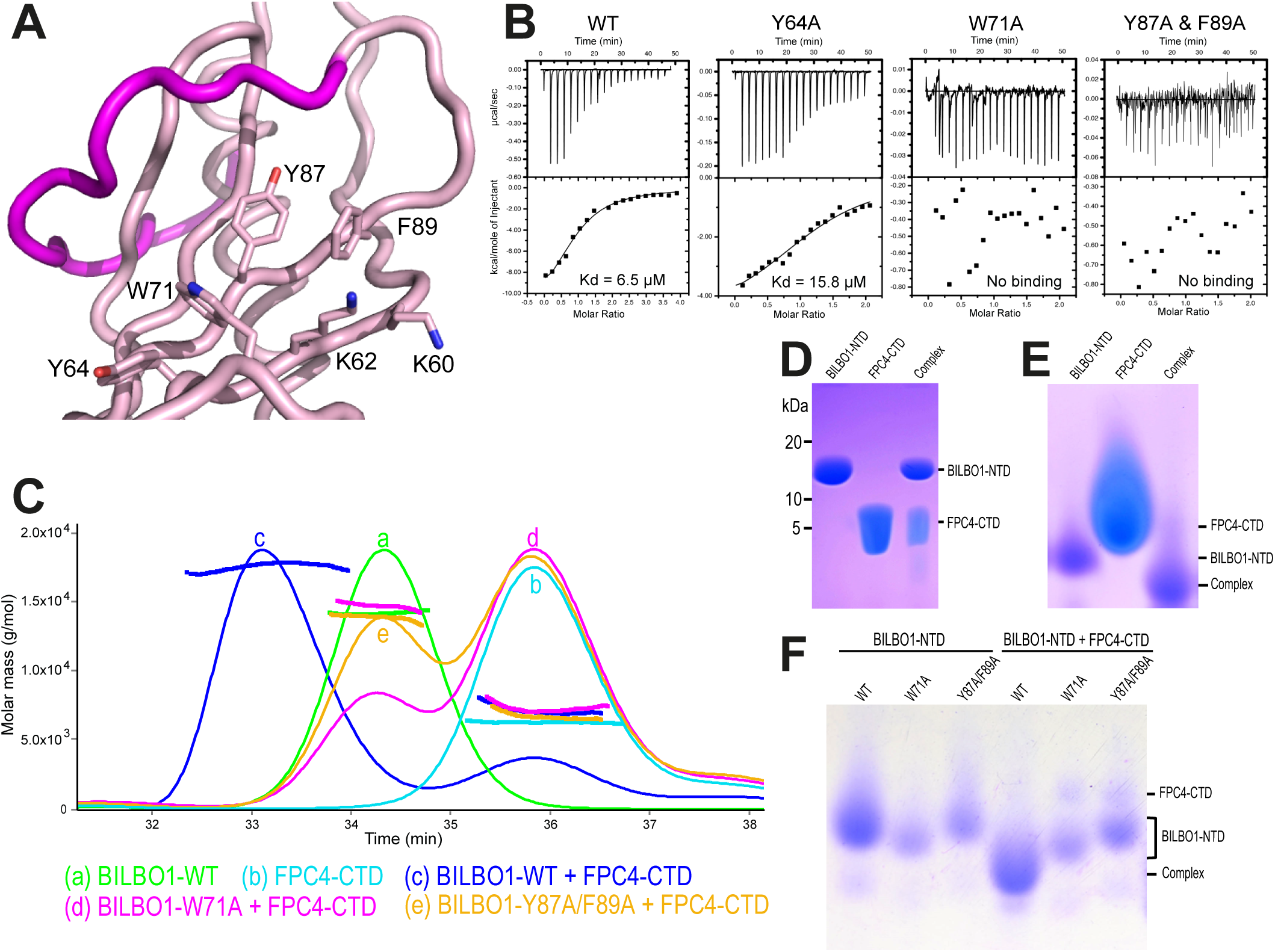
The aromatic residues of the TbBILBO1-NTD pocket are required for TbFPC4 binding. *A*, Cartoon view of the hydrophobic pocket of TbBILBO1-NTD with all mutated residues shown in sticks and labeled. *B*, ITC results of wild-type (WT) and mutants of TbBILBO1-NTD with TbFPC4-CTD. Mutants W71A and Y87A/F89A showed no interaction, whereas mutants Y64A and K60A/K62A showed slightly reduced binding affinity. *C*, SLS profiles of individual TbBILBO1-NTD (a) and TbFPC4-CTD (b), and TbFPC4-CTD mixed with WT (c) and mutants of TbBILBO1-NTD (d and e). The highest peaks in the elution profiles are normalized to the same height for easy comparison. *D*, Purified proteins of TbBILBO1-NTD, TbFPC4-CTD, and their complex from SEC on an SDS-PAGE gel and visualized by Coomassie staining. *E*, The same samples as in *D* on a native gel. *F*, Comparison of WT and mutants of TbBILBO1-NTD alone or in complex with TbFPC4-CTD on a native gel. In contrast to the stable complex formed between the WT TbBILBO1-NTD and TbFPC4-CTD, mutants W71A and Y87A/F89A did not form complexes with TbFPC4-CTD.

Interaction tests by ITC experiments revealed that the W71A mutant completely abolished the interaction of TbBILBO1-NTD with TbFPC4, whereas Y64A only reduced the binding affinity by approximately 2.5 fold (Fig. 4B). To further validate the dramatic effect of those mutants that abolish the interaction, we additionally carried out SEC/SLS analysis using purified recombinant proteins. The results showed that, in contrast to the stably formed binary complex between wild-type TbBILBO1-NTD and TbFPC4-CTD, neither of the two TbBILBO1 mutants, W71A and Y87A/F89A, could form a stable complex with TbFPC4 (Fig. 4C), which further suggests that these three aromatic residues play pivotal roles in the interaction between the two proteins.

Furthermore, we checked the interaction of TbBILBO1 and TbFPC4 using EMSA on native gels. As expected, the purified TbBILBO1/TbFPC4 complex showed a clearly shifted band comparing to the bands of the two individual proteins (Fig. 4D & E). Similarly, the mixture of TbBILBO1-NTD and TbFPC4-CTD in an equimolar ratio also showed a shifted band corresponding to the formed complex (Fig. 4F). In contrast, mixing TbFPC4-CTD with TbBILBO1-NTD mutants W71A or Y87A/F89A resulted in two separate bands of the two individual proteins (Fig. 4F).

### Influence of the TbBILBO1 mutants on TbBILBO1-TbFPC4 interaction in U-2 OS cells

It was previously shown that the strong interaction between TbBILBO1 and TbFPC4 drives the colocalization of the two proteins when ectopically expressed in mammalian U-2 OS cells (8). We carried out a similar assay to further check the effect of the mutations on the interaction between TbBILBO1 and TbFPC4 under *in vivo* conditions. Similarly to the previous report, expression of TbBILBO1 alone in U-2 OS cells generated filamentous polymers (Fig. 5A), while expression of GFP-tagged TbFPC4 showed that it decorated a filamentous network that matched the previously determined colocalization with microtubules (Fig. 5B). When co-expressed with TbBILBO1, TbFPC4 was only associated with the TbBILBO1 polymers (Fig. 5C). Co-expression of TbBILBO1-Y64A and TbFPC4 showed that TbFPC4 is associated with the polymers formed by TbBILBO1-Y64A, but some microtubule interaction was also observed (Fig. 5D, inset), suggesting a decrease in the binding affinity to TbBILBO1. When co-expressed with TbBILBO1-W71A, TbFPC4 was not associated with the TbBILBO1 polymers but gave a microtubule pattern, demonstrating a complete absence of binding to the TbBILBO1-W71A form (Fig. 5E), which was consistent with the *in vitro* data (Fig. 4).

**FIGURE 5.**
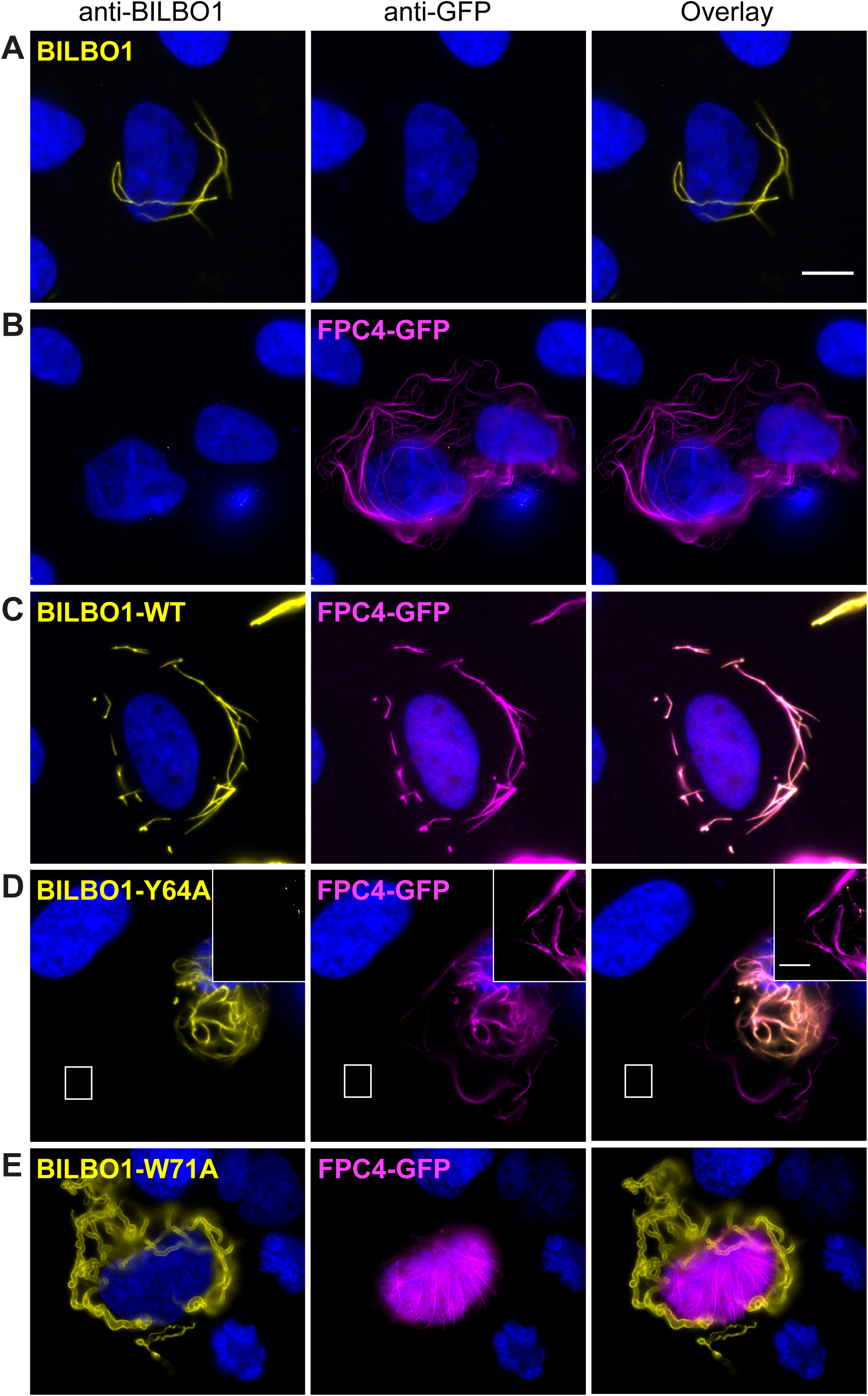
The aromatic residues of the TbBILBO1-NTD pocket are required for TbFPC4 recruitment *in vivo*. *A*, Expression of TbBILBO1 alone in U-2 OS cells generated filamentous polymers. *B*, Expression of TbFPC4-GFP alone showed its tight association with microtubules. *C*, Co-expressed TbBILBO1 and TbFPC4-GFP proteins colocalized with each other. *D*, Co-expression of TbBILBO1-Y64A and TbFPC4-GFP showed relatively robust overlap of TbFPC4 with the TbBILBO1-Y64A polymers, but some microtubule interaction is also observed (Inset). *E*, Co-expression of TbBILBO1-W71A with TbFPC4-GFP showed no colocalization of the two proteins; TbFPC4 showed microtubule pattern. Scale bars represent 10 μm, and 1 μm in the zoom images.

### Effect of the TbBILBO1 mutants in trypanosomes

We assessed the effect of the tetracycline-inducible ectopic expression of the Ty1 epitope-tagged TbBILBO1, Ty1 epitope-tagged Y64A and W71A TbBILBO1 mutants, as well as another mutant D65K/E66K/E67K (DEE/KKK), in procyclic (PCF) trypanosomes. The mutant DEE/KKK was tested because these three residues are not only highly conserved but also next to the peripheral aromatic residue Y64.

We used Western blotting to assess the expression level of Ty1-tagged TbBILBO1 (Fig. 6A), and cell growth to assess phenotypes (Fig. 6B-E). Overall, induction of the expression of wild-type _Ty1_TbBILBO1 and of mutant Y64A had no or little impact on cell growth (Fig. 6B, C), suggesting that Y64 is not an essential residue for BILBO1 function. However, after 24 hours of induction, expression of _Ty1_TbBILBO1-W71A resulted in a dominant-negative lethal phenotype (Fig. 6D), demonstrating that this residue is essential for TbBILBO1 function in the parasite. Interestingly, despite the close location of D65/E66/E67 to the hydrophobic pocket, there was no observable effect of TbBILBO1-DDD/KKK on cell growth at all (Fig. 6E).

**FIGURE 6.**
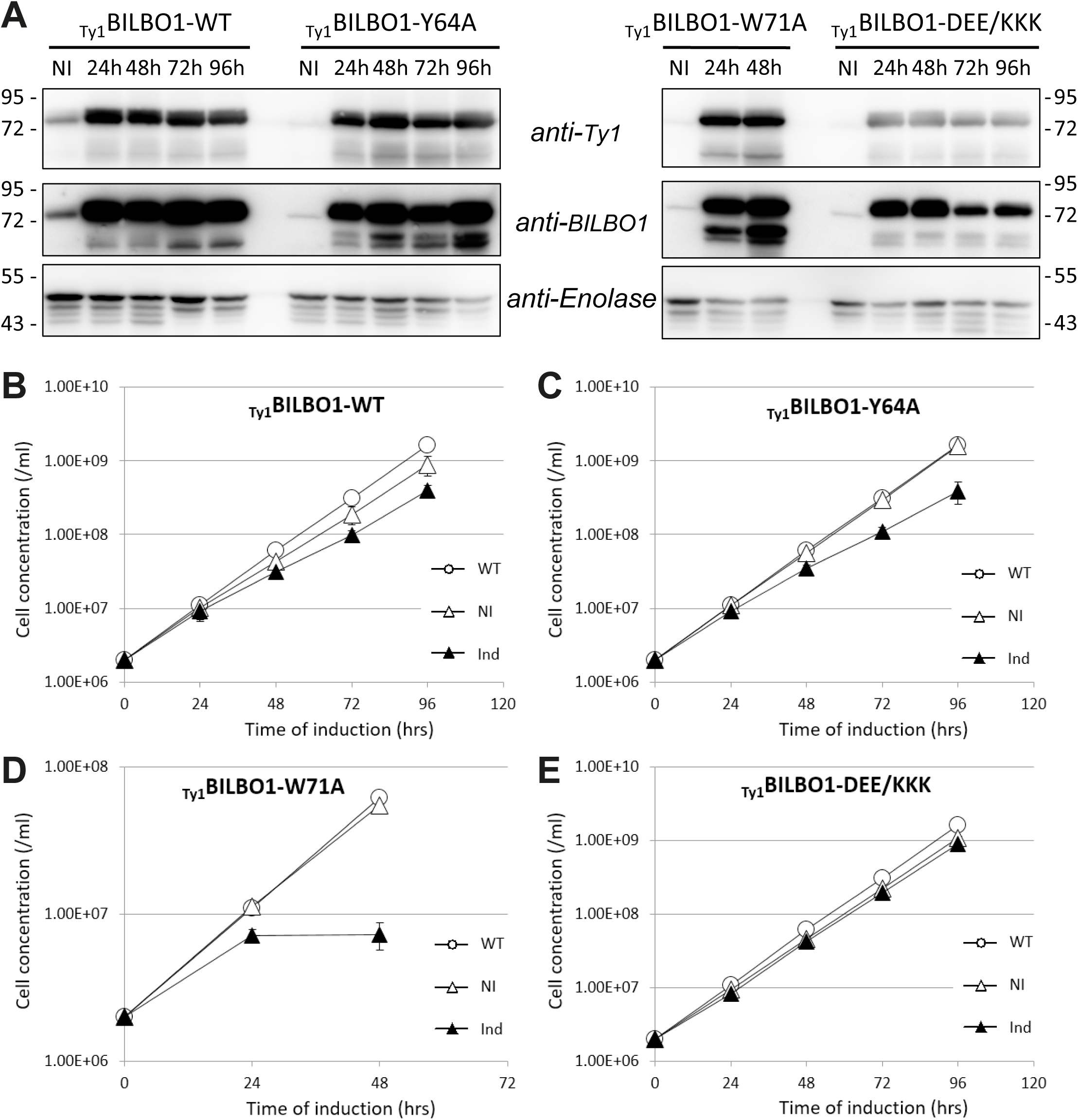
Residue W71 on TbBILBO1 is essential for its function in *T. brucei*. *A*, WB for the PCF *T. brucei* cells, non-induced (NI) or induced up to 96 hours for the expression of Ty1-tagged TbBILBO1-WT, TbBILBO1-Y64A, TbBILBO1-W71A, or TbBILBO1-DEE/KKK. Anti-Enolase was used as a loading control. *B-E*, Growth curves for the PCF *T. brucei* cells from the same experiments as in *A*. The error bars represent the standard error from three independent experiments.

## DISCUSSION

BILBO1 was the first identified component of the FPC (5). It consists of four structural domains, including the globular NTD that was shown to be essential for the function of TbBILBO1 in the cell (10). We have previously reported the NMR structure of TbBILBO1-NTD, which demonstrated that it adopts a ubiquitin-like fold (10). The 1.6-Å resolution crystal structure of TbBILBO1-NTD reported here shows an overall similar conformation to the NMR structure, with an extended β sheet formed by five strands and a long α helix packed diagonally across the surface of the β sheet (Fig. 1).

In the NMR structure, the C-terminal region of TbBILBO1-NTD, spanning residues 101-110, was poorly defined. Surprisingly, in the newly determined crystal structure this region was rigidly bound to the structural core of the protein. How can we explain the discrepancy observed between these two structures?

Firstly, the construct used for NMR studies encoded residues 1-110, which was designed based on the conservation analysis and *a priori* structural prediction. Sequence alignment of BILBO1 homologs from various kinetoplastid species showed a clear boundary at the residue P110 between the highly-conserved NTD and the following variable linker (10). Furthermore, secondary structure prediction suggested that there were no helices or strands after residue 90. Therefore, we considered only residues 1-110 of TbBILBO1 in our original structural studies, which was much shorter than the protein we later used for crystallization reported in this study, i.e. residues 1-175.

During our NMR studies, we were able to determine the core part of the structure up to residue Q100. However, no long-range NOE constraints could be confidently assigned for residues 101-110. Notably, there are four proline residues (P104, P105, P109, & P110) in that region (Fig. 2A), which could not be assigned due to their lack of backbone amide hydrogens. These were probably the main reasons for the undefined C-terminal loop in the NMR structure.

Second, the crystal structure was solved based on the naturally degraded product of a longer construct of TbBILBO1 covering residues 1-175 (11). The extra residues in the crystallization construct might have played a role in stabilizing the loop to make it bind more tightly to the structural core of TbBILBO1. Indeed, there were multiple hydrophobic interactions as well as water-mediated hydrogen bonds between residues 110-114 and the C-terminal part of the α-helix in the crystal structure of TbBILBO1-NTD (Fig. 2D). Notably, these interactions are mediated mostly by the backbone atoms of these residues, which explains why they are not conserved in the primary sequence, but could yet still participate in the binding of the loop to the structural core of TbBILBO1-NTD.

The core of the TbBILBO1-NTD structure (residues 1-91) is topologically identical to the ubiquitin fold (Supplementary Fig. 1). However, it was striking to observe such an enormously long rigid loop in the crystal structure, particularly for such a small compact domain. The loop has a stable conformation as demonstrated by the well-defined electron densities and comparable temperature factors of all the loop residues (Fig. 2B and C). Further check of molecular packing in the crystal structure demonstrated the majority of the loop is situated in a cavity without contacting the neighboring molecules (Supplementary Fig. 2). All these evidences strongly suggest that the long loop in the crystal structure is in its native state. The highly curved conformation of the loop is critical for the formation of the unique horseshoe-like hydrophobic pocket (Fig. 3), which turned out to be the binding site of TbFPC4 (Fig. 4).

Although we did not know the binding partner of TbBILBO1-NTD when we reported the NMR structure, our mutagenesis studies demonstrated that it contains a surface patch critical for TbBILBO1 function *in vivo*. Compared to the NMR structure, the crystal structure reported here not only provides a higher resolution structure of TbBILBO1-NTD, but also sheds light on the horseshoe-like conformation of the hydrophobic pocket, which was shown to be relatively flat and irregular in the previous NMR structure due to the poorly defined C-terminal loop as discussed above.

Regarding the binding of TbFPC4 to TbBILBO1-NTD, our *in vitro* binding experiments demonstrated that mutating the three central aromatic residues, Y87, F89 and W71, completely abolished the interaction of TbBILBO1-NTD with TbFPC4, which suggests that TbFPC4 directly contacts these aromatic residues (Fig. 4, Fig. 5). *In vivo* examination of these mutants revealed that the three central aromatic residues play essential roles in controlling cell growth (Fig. 6), which is consistent with the previously reported results (8,10), and further confirms the importance of the conserved hydrophobic pocket.

In our mutagenesis studies we found that residue W71, which gates the entrance to the horseshoe-like pocket on TbBILBO1-NTD, is absolutely required for both TbFPC4 binding (Fig. 4B, Fig. 5E) and cell growth (Fig. 6D). However, mutating Y64, the residue outside the horseshoe-like pocket, only mildly reduced the binding of TbBILBO1-NTD to TbFPC4 (Fig. 4B; Fig. 5D). Further, the Y64A mutant did not cause dramatic effect in cell growth when overexpressed *in vivo* (Fig. 6C). We therefore believe that the corresponding interacting region on TbFPC4-CTD, which is predicted to overall lack any secondary structures and may thus adopt an extended conformation, likely binds horizontally across the β sheet to allow it pass through the gap of the horseshoe-like pocket (Fig. 7). Nevertheless, the binding site of TbFPC4 on TbBILBO1-NTD seems to be limited to the hydrophobic pocket and neighboring residues in the close proximity, as mutations of the three negatively charged residues, D65, E66, and E67, which are slightly farther away from the pocket, did not affect either the intermolecular interaction or cell growth (Fig. 6E).

**FIGURE 7.**
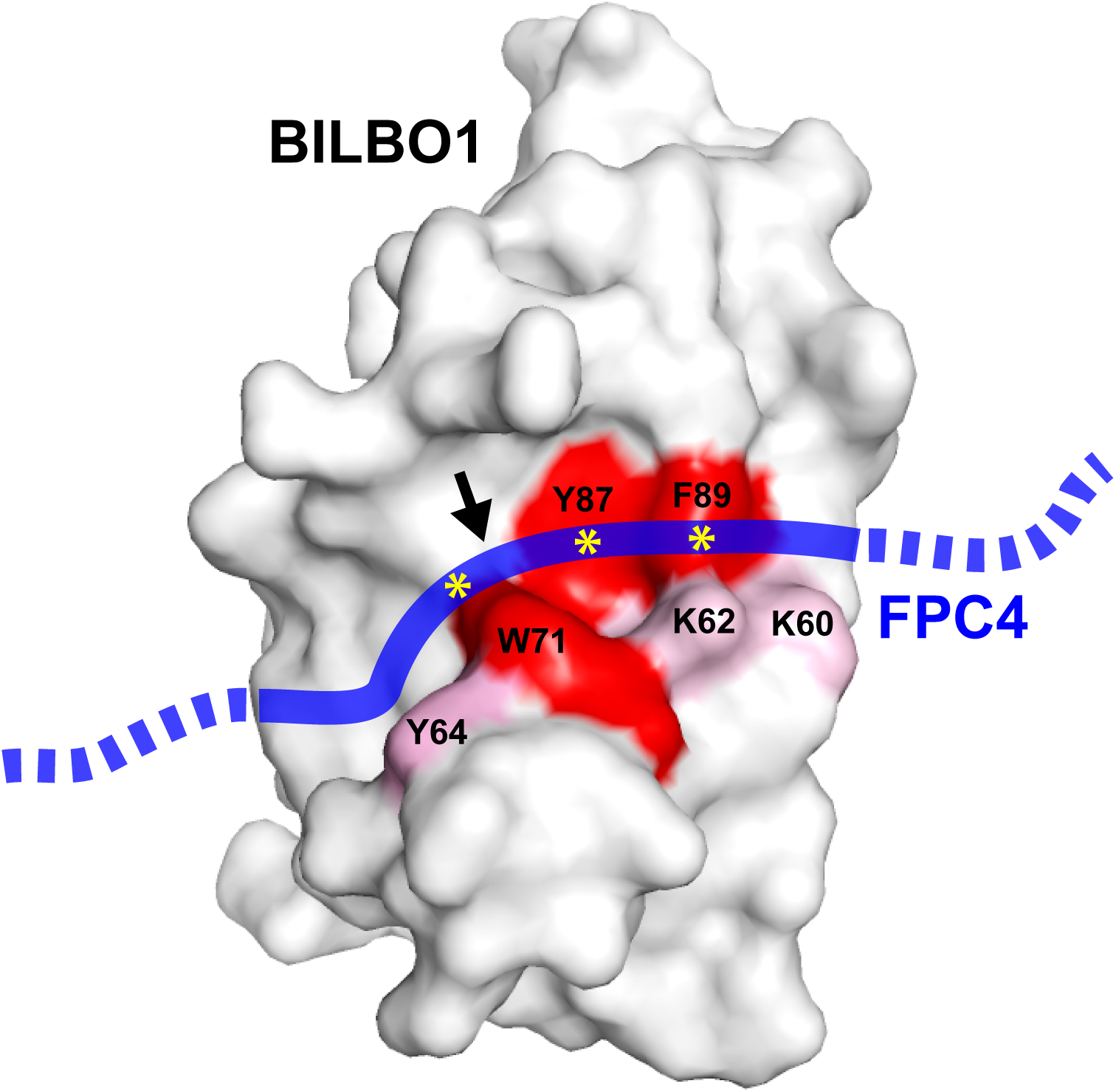
A hypothetical model depicting the interaction between TbBILBO1-NTD and TbFPC4-CTD. TbBILBO1-NTD is shown in a gey surface plot with residues essential for TbFPC4 binding highlighted in red, and those partially affecting TbFPC4 binding colored in pink. TbFPC4 is depicted as a blue line, with regions involved in critical interactions marked as asterisks. The arrow points to the part in TbFPC4 that passes through the gap of the horseshoe-like pocket to reach the peripheral binding site outside of the pocket.

The close correlation of the hydrophobic pocket in mediating the interaction of TbBILBO1 with TbFPC4 and its requirement for normal cell growth suggests that the main function of TbBILBO1-NTD might be to act in concert with TbFPC4 to regulate the structural and/or function of the FPC in trypanosomes. Previous studies have shown that an N-terminal region of TbFPC4 binds microtubules, which prompted the proposal that TbFPC4 might serve as a linker to connect the FPC to the microtubule quartet (8).

In summary, we have reported a high resolution crystal structure of TbBILBO1-NTD (residues 1-115), which reveals an unusual rigid loop critical for defining the horseshoe-like hydrophobic pocket responsible for the binding of its partner, TbFPC4. Notably, by now there have been no reports about a similar surface pocket on any known ubiquitin-like fold containing proteins except for TbBILBO1-NTD. Future studies will try to map the exact region on TbFPC4 and find out the molecular detail of their interaction.

Given the relatively low binding affinity between TbBILBO1 and TbFPC4, with a Kd of ~5 μM, it is probably worth checking whether a modified synthetic polypeptide with significantly higher affinity than the native sequence of TbFPC4 could be found, which could be potentially used as an inhibitor to disrupt the interaction between TbFPC4 and TbBILBO1 in the parasite. Alternatively, screening small molecular libraries may be carried out to identify candidate inhibitors that are tightly bound to the hydrophobic pocket to prevent the formation of the complex between TbBILBO1-NTD and TbFPC4.

## CONFLICT OF INTEREST

The authors declare that they have no conflict of interest.

## ACKNOWLEDGEMENT

* This work was supported by funding from the Max Perutz Labs and grants P24383-B21 and P28231 from the Austrian Science Fund (FWF) to GD, the Bordeaux University and the Centre National de la Recherche Scientifique (CNRS) to DRR and MB, and the LabEx ParaFrap (ANR-11-LABX-0024) to DRR. YP has been supported during 2016-2019 by the “Integrative Structural Biology” PhD program (W-1258 Doktoratskollegs) funded by the FWF. KV was supported by an OeAD graduate scholarship during 2009-2012. NA was supported by a postdoctoral fellowship provided by the Austrian Agency for International Cooperation in Education & Research (OeAD-GmbH) during January – September 2019. We thank Frédéric Bringaud (Bordeaux University) for the anti-Enolase antibody and P. Bastin (Pasteur Institute) for the anti-Ty1 (BB2) antibody. We also greatly appreciate Dr. Brooke Morriswood for critical reading of the manuscript.

## Notes

#### Summary of Updates

We have substantially reworked our manuscript, addressing all conceptual comments, clarifying confusions in the text, and making modifications wherever are necessary. Following are the four major changes we have made in the revised version. 1. New Supplementary Figure 1 is added to show the Dali search results, together with a new paragraph in the Results section to summarize the findings. 2. New Supplementary Figure 2 is added to show crystal packing and the neighboring environment of the C-terminal loop in the TbBILBO1-NTD. 3. Fig. 2B is revised to show the electron density map of the loop. 4. Fig. 4C is updated to provide a high resolution image of the SLS data.

