## Supplemental Fig. 1 & 2 for "Crystal structure of the TbBILBO1 N-terminal domain reveals a ubiquitin fold with a long rigid loop for the binding of its partner"

**LEGENDS for SUPPLEMENTARY FIGURES**

SUPPLEMENTARY FIGURE 1. **TbBILBO1-NTD structure consists of a ubiquitin-like core with a unique long C-terminal loop.** *A*, Correlation matrix plot shows the correlation of DALI-matched 25 top ranking homologous structures along positions of the TbBILBO1-NTD sequence (22). Residues 3-91 show strong positive correlation, which indicates a conserved conformation for all these proteins. The C-terminal loop of TbBILBO1-NTD (residues 92-115) shows very weak correlation, suggesting that such loop is not conserved in other ubiquitin-like structures. The x-axis shows PDB residue numbers and the y-axis labels sequential residue numbers. *B*, Two orthogonal views of superposition of TbBILBO1-NTD (light green) with the top 11 matched structures found by the DALI server (22) that have a Z-score ≥ 7.0 and r.m.s.d. ≤ 2.3 Å. Residue P92 of TbBILBO1 that is located at the junction of the conserved ubiquitin domain and C-terminal loop is marked.

SUPPLEMENTARY FIGURE 2. **The rigid conformation of the loop is in its native state, rather than a crystallization artifact.** *A*, Crystal packing around the C-terminal loop with two orthogonal views. Residues 96-115 of the loop is shown as sticks in magenta. *B*, 2Fo-Fc maps (1.5 σ) with a top view looking down onto the plan of the curved loop. The whole loop is situated mostly in a cavity with only its C-terminal part contacting a neighboring molecule.

Supplementary Fig. 1

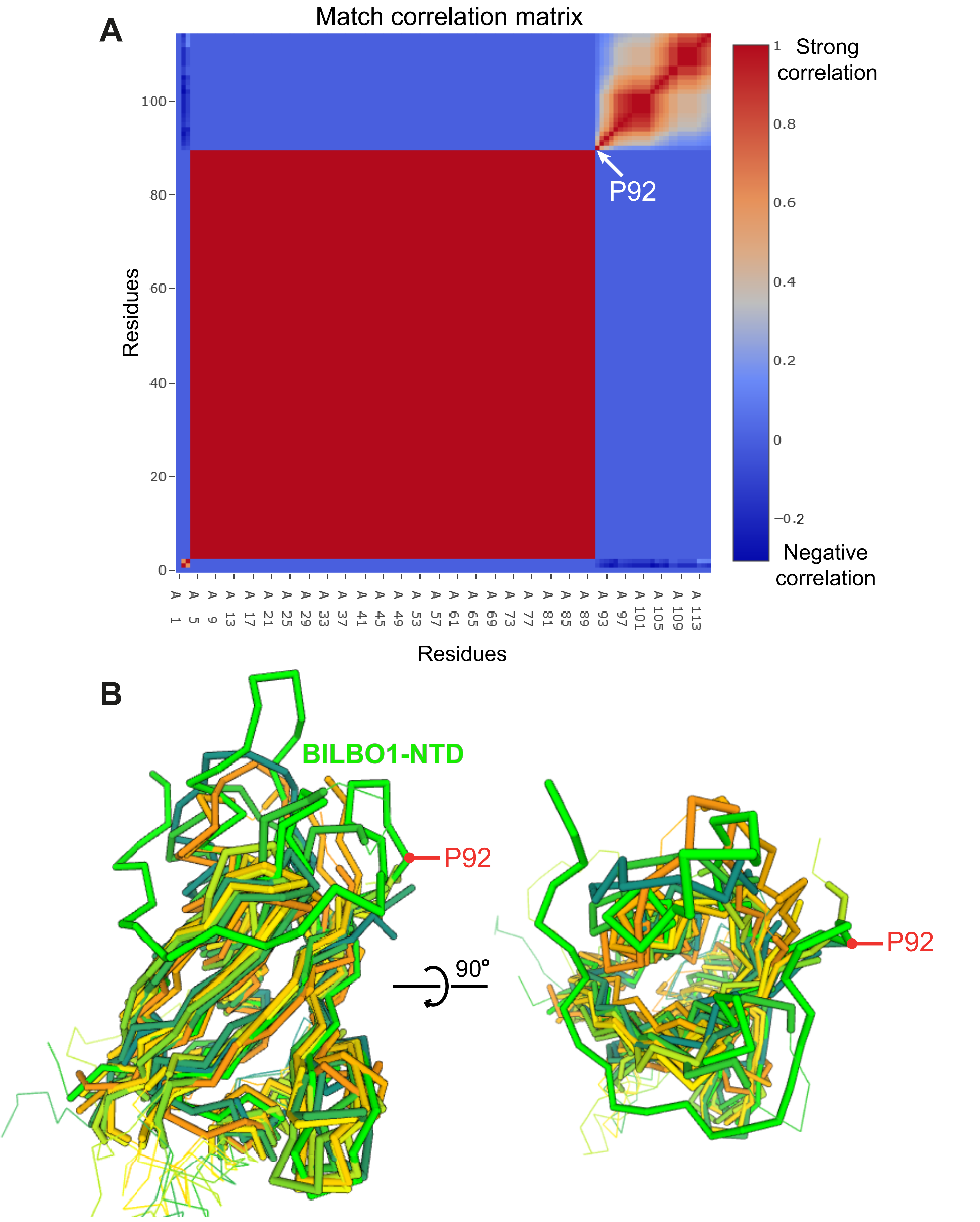

Supplementary Fig. 2

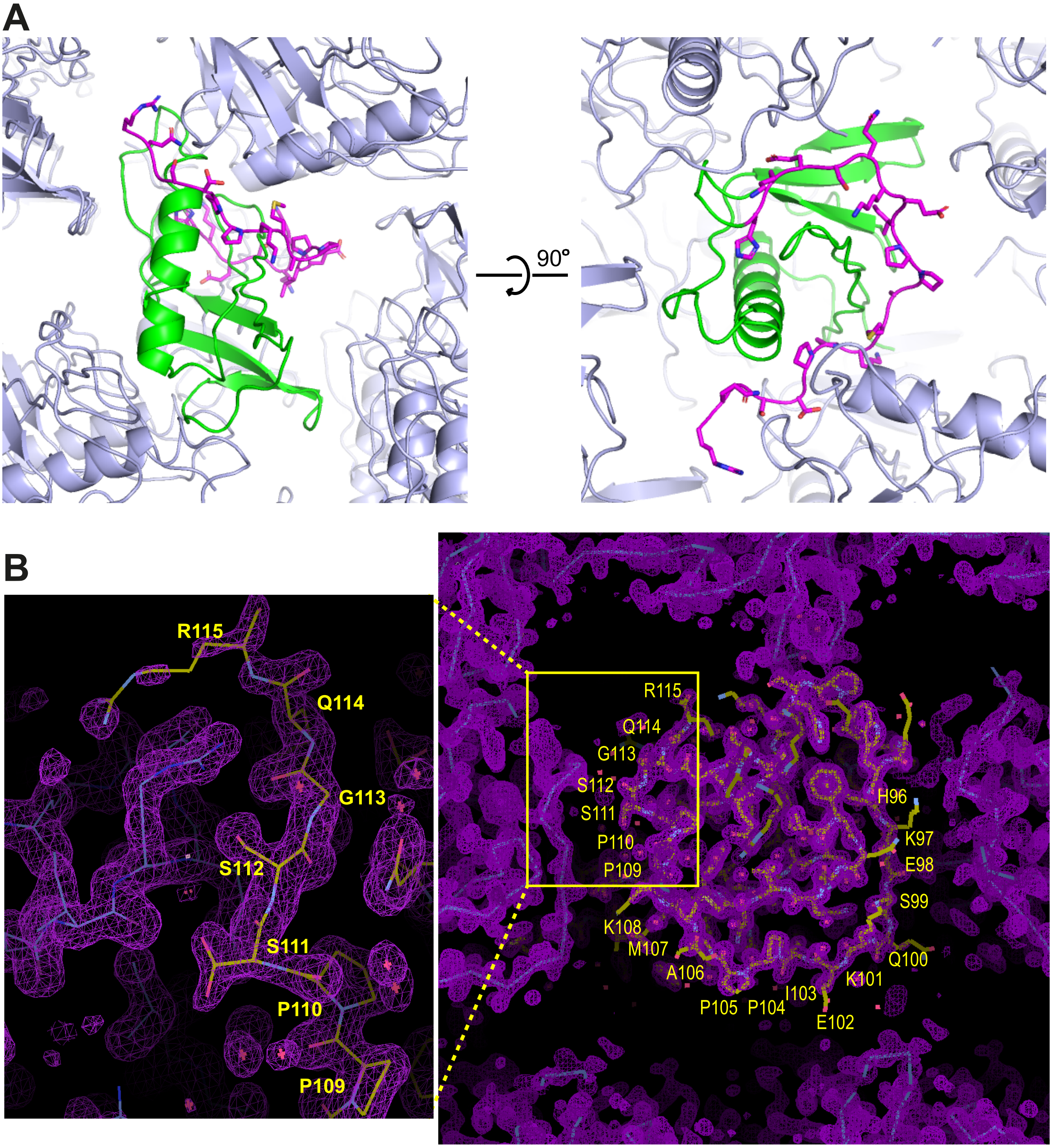
